# CRISPR-Cas9 mediated genome editing in vancomycin resistant *Enterococcus faecium*

**DOI:** 10.1101/745786

**Authors:** Vincent de Maat, Paul B. Stege, Mark Dedden, Maud Hamer, Jan-Peter van Pijkeren, Rob J.L. Willems, Willem van Schaik

## Abstract

The Gram-positive bacterium *Enterococcus faecium* is becoming increasingly prevalent as a cause of hospital-acquired, antibiotic-resistant infections. There is thus an urgent need to mechanistically characterize the traits that contribute to the emergence of *E. faecium* as a multidrug resistant opportunistic pathogen. A fundamental part of research into *E. faecium* biology relies on the ability to generate targeted mutants, but this process is currently labour-intensive and time-consuming, taking 4 to 5 weeks per mutant. In this report we describe a method relying on the high recombination rates of *E. faecium* and the application of the Clustered Regularly Interspaced Short Palindromic Repeat (CRISPR)-Cas9 genome editing tool to more efficiently generate targeted mutants in the *E. faecium* chromosome. Using this tool and the multi-drug resistant clinical *E. faecium* strain E745, we generated a deletion mutant in the *lacL* gene, which encodes the large subunit of the *E. faecium* β-galactosidase. Blue/white screening using 5-bromo-4-chloro-3-indolyl-β-D-galactopyranoside (X-gal) could be used to distinguish between the wild-type and *lacL* deletion mutant. We also inserted two copies of *gfp* into the intrinsic *E. faecium* macrolide resistance gene *msrC* to generate stable green fluorescent cells. We conclude that CRISPR-Cas9 can be used to generate targeted genome modifications in *E. faecium* in 3 weeks, with limited hands-on time. This method can potentially be implemented in other Gram-positive bacteria with high intrinsic recombination rates.

## Introduction

Microbial antibiotic resistance is currently recognised as a global threat to human health (Ferri *et al.* 2017). Enterococci are among the most problematic multi-drug resistant bacteria causing infections among hospitalised patients, contributing to 10,000 to 25,000 deaths per year in the USA alone (McKinnell *et al.* 2012). Clinically, the two most important enterococcal species are *Enterococcus faecalis* and *Enterococcus faecium*. While historically *E. faecalis* has been the most prominent enterococcal pathogen, since the 1990s *E. faecium* has rapidly emerged as a nosocomial pathogen of major importance. Infections caused by *E. faecium* are generally more difficult to treat as vancomycin resistance is more widespread in *E. faecium* than in *E. faecalis* (Gilmore, Lebreton and Schaik 2013; García-Solache and Rice 2019). Until we understand the molecular underpinnings that contribute to the transfer of antibiotic-resistant genes, and pathogenicity, we will be hampered in our ability to develop treatment strategies. To drive functional studies, efficient genome editing tools are essential, which are currently lacking. Current methods to generate targeted mutations in *E. faecium* mostly rely on allelic exchange between the chromosome and a suicide vector which contains an antibiotic resistance cassette and sequences that flank the target site on the *E. faecium* genome (Maguin *et al.* 1996; Nallapareddy, Singh and Murray 2006; Zhang *et al.* 2012). The antibiotic cassette can be removed using the Cre-*lox* system, but a single *lox* site remains as a scar (Zhang *et al.* 2012). These protocols are time-consuming, taking upwards of 4 to 5 weeks. The process involves several days of sub-culturing and selection of colonies on media with different antibiotics, to screen for a double cross-over event and then removal of the resistance marker by Cre-*lox*. In addition, extensive screening by colony PCR is needed to retrieve the desired mutant. The process to generate targeted mutants in *E. faecium* was improved by the use of counter-selection system against single cross-over mutants by the use of *pheS*\*, a mutated allele of the *E. faecalis* phenylalanyl tRNA synthetase α-subunit that confers susceptibility to p-chloro-phenylalanine in enterococci (Kristich, Chandler and Dunny 2007; Thurlow, Thomas and Hancock 2009; Somarajan *et al.* 2014; Bhardwaj, Ziegler and Palmer 2016).

To further expand the genetic toolbox for multi-drug resistant *E. faecium*, we explored the use of clustered regularly interspaced palindromic repeats (CRISPR) and its associated Cas9 protein to generate mutants in *E. faecium*. The Cas9 nuclease introduces double-stranded breaks in DNA that is targeted by a CRISPR and, together with other CRISPR-asssociated proteins, serves as a defence against invading bacteriophages in prokaryotes (Brouns *et al.* 2008). The combination of CRISPR and Cas9 has been successfully used for genome editing in eukaryotes where CRISPR-Cas9 drives the generation of mutants by inducing double-stranded DNA breaks which are then repaired by non-homologous end-joining (NHEJ) (Cong *et al.* 2013). While NHEJ systems are present in some bacteria (Shuman and Glickman 2007), most prokaryotes can only escape the lethal effect of CRISPR-Cas9 targeting a chromosomal site by utilising homologous recombination (HR). One approach to use CRISPR-Cas to identify recombinant genotypes is to introduce a vector that contains DNA identical to the flanking sequence of the target region while the cell produces Cas9 and a CRISPR-array homologous to the target sequence. Most surviving cells will have undergone a HR event thereby escaping CRISPR-Cas mediated killing. (Jiang *et al.* 2013; Wang *et al.* 2015, 2018). Genome editing approaches using HR and CRISPR-Cas9 have been used for numerous bacterial species, including Gram-positive lactic acid bacteria (Mougiakos *et al.* 2016; Leenay *et al.* 2019).

In this study, we aimed to develop a CRISPR-Cas9 based genome editing approach for *Enterococcus faecium*. We adapted a CRISPR-Cas9 based genome editing approach previously developed for the lactic acid bacterium *Lactobacillus reuteri* (Oh and Van Pijkeren 2014), relying on the high intrinsic recombination rate of *E. faecium* for allelic exchange combined with CRISPR-Cas9 to counterselect against wild-type cells.

## Materials and methods

### Bacterial strains, plasmids, growth conditions, and oligonucleotides

The vancomycin-resistant *E. faecium* strain E745 (Zhang *et al.* 2017) was used throughout this study. This strain was isolated from a rectal swab of a hospitalized patient, during routine surveillance of a VRE outbreak in a Dutch hospital. Unless otherwise mentioned, *E. faecium* was grown in brain heart infusion broth (BHI; Oxoid) at 37 °C. The *E. coli* strain EC1000 (Leenhouts *et al.* 1996) was grown in Luria-Bertani (LB) medium at 37°C while shaking at 200 rpm. *L. lactis* MG1363 was grown in M17 broth supplemented with 0.5% w/v lactose. When required, antibiotics were used at the following concentrations: erythromycin 50 μg ml^−1^ for *E. faecium* and 5 μg ml^−1^ for *L. lactis* and spectinomycin 200 μg ml^−1^ for *E. faecium*, 100 μg ml^−1^ for *E. coli*, and tetracycline 10 μg ml^−1^ for *L. lactis*. Where indicated, plates were supplemented with 0.2% 5-bromo-4-chloro-3-indolyl-β-D-galactopyranoside (X-gal). The vectors pREG696 (Grady and Hayes 2003), pWS3 (Zhang *et al.* 2011), and pET-3α (Novagen) were obtained from our laboratory’s culture collection. pREG696-*gfp* was derived from pREG696 by inserting the *gfp* gene under the control of the promoter of the *bacA* gene (pBac) of *E. faecalis* (Heikens, Bonten and Willems 2007) in the NotI and XhoI restriction sites of pREG696 (J. Top, personal communication). Plasmids pVPL3004 and pVPL3115 were previously described in (Oh and van Pijkeren, 2014). The sequences of the oligonucleotides used in this study are listed in Table 1.

**Table 1:**
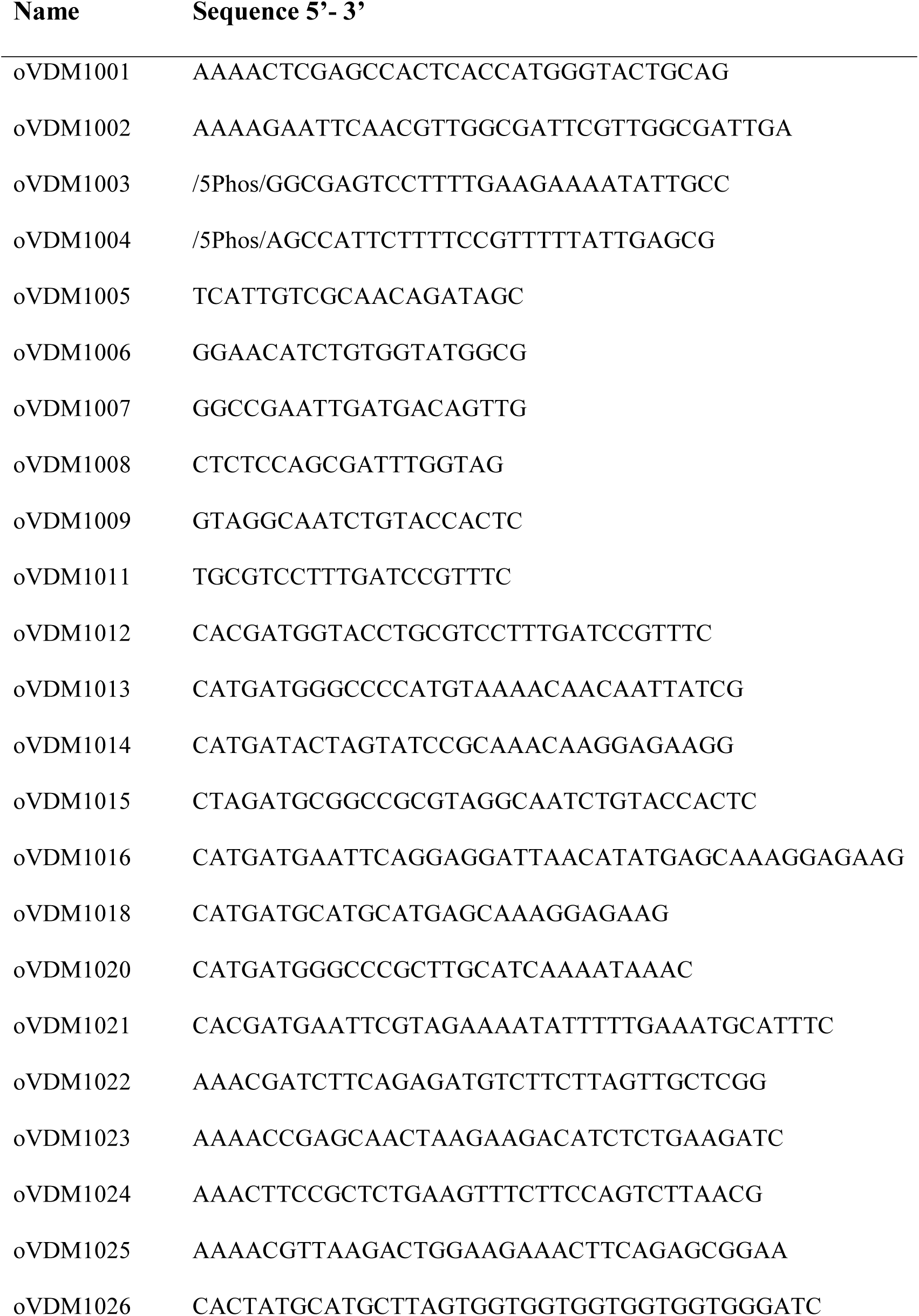
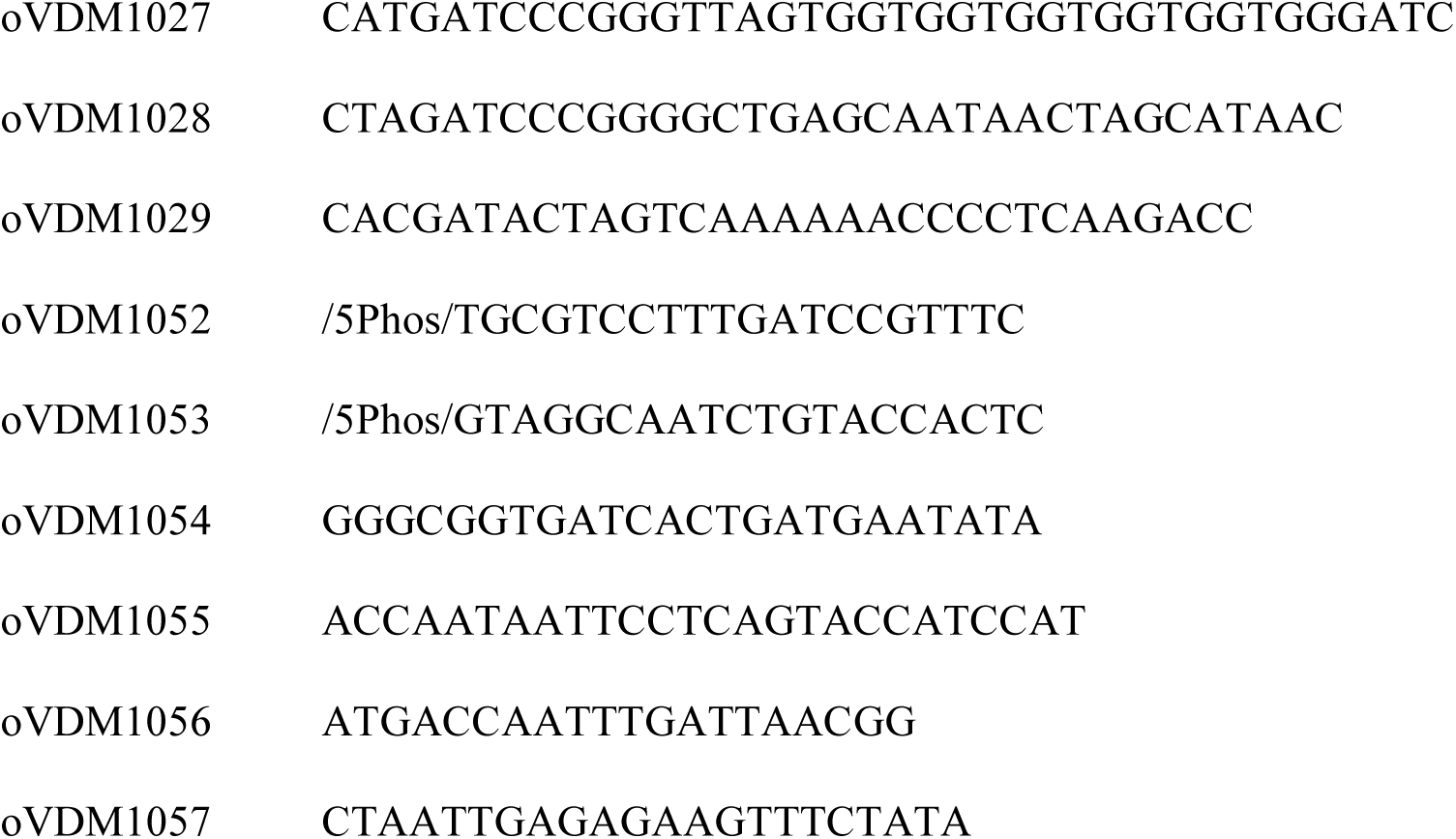
List of oligonucleotides used in this study.

### Isolation and transformation of plasmids

Plasmid isolation from *E. coli* was performed using the GeneJET plasmid miniprep kit (Thermo Fischer Scientific, Bleiswijk, the Netherlands) according to the manufacturer’s instructions. Isolation of plasmids from *L. lactis* was as described previously (O’Sullivan and Klaenhammer 1993) with slight modifications. In short, 5 ml overnight cultures were pelleted by 10 minute centrifugation at 3000 g. The pellet was resuspended in 250 μl THMS-buffer (30 mM Tris-HCL pH 8, 25% sucrose, 3 mM MgCl_2_) supplemented with 2 mg ml^−1^ lysozyme. The cell suspension was incubated for 10 minutes at 37°C after which 500 μl 1% SDS in 0.2 M NaOH was added. The tubes were mixed gently and incubated on ice for 5 minutes. 375 μl ice-cold 3M potassium acetate pH 5.5 was added and the mixture was mixed by inversion, followed by incubation on ice for 5 minutes. Cell debris was pelleted via centrifugation at 20,000 g for 5 min, after which the supernatant was transferred to a new tube and an equal amount of isopropanol was added. After 10-minute incubation at room temperature the tubes were centrifuged at 20,000 g for 10 min to precipitate the DNA. The pellet was washed with 70% ethanol, air dried, and dissolved in sterile dH_2_O. Transformation of plasmids into *E. faecium* E745 was performed as previously described (Zhang *et al.* 2012).

### Construction of the pVDM1001 CRISPR delivery vector and generation of *lacL-*deletion and *gfp*-insertion mutants

We first aimed to construct a vector that could be used for genome editing in *E. faecium* E745. This vector, termed pVDM1001, was created by cloning a 0.7-kbp fragment which contains the CRISPR sequences from pVPL3115 in the XhoI and EcoRI sites of pWS3. The fragment was amplified from pVPL3115 using the primers oVDM1001 - oVDM1002. The pVDM1001 vector was then implemented for the generation of a *lacL* deletion and *gfp* insertion mutant by modifying the CRISPR sequence via digestion with BsaI and annealing two oligos, oVDM1022-oVDM1023 and oVDM1024-oVDM1025, which contain a protospacer targeting *lacL* or *msrC,* respectively. This created pVDM-x*lacL* and pVDM-x*msrC*. Next a DNA template consisting of a 365bp upstream region of *lacL* fused together with a 225 bp downstream region of *lacL* was ordered. (Table S1) and amplified using oVDM1003 - oVDM1004. The amplified template was cloned into pVDM-x*lacL* after digestion with SmaI and a blunt end ligation creating pVDM-*ΔlacL*.

To create a *gfp* knock-in construct we amplified 773 bp upstream region of *msrC* and a 507 bp fragment overlapping with the 3’ region of *msrC* using primers oVDM1012 - oVDM1013 and oVDM1014 - oVDM1015, respectively. Each fragment was separately cloned into pWS3 using KpnI-ApaI for the upstream fragment and SmaI-NotI for the downstream fragment, creating pWS3-*msrCup* and pWS3-*msrCdwn*, respectively. Downstream of the *msrCup* fragment a pBac promotor was inserted. The promotor site was amplified from pREG696-*gfp* using primers oVDM1020 – oVDM1021 and inserted after ApaI-EcoRI digestion creating pWS3-*msrCup*-pBac. To pWS3-*msrCdwn* a T7 terminator was added which was amplified from pET3α using primers oVDM1028-oVDM1029 and digested with SmaI-SpeI to create pWS3-T7-*msrCdwn*. pWS3-*msrCup*-pBac was then digested with KpnI-EcoRI and the *msrCup-*pBac fragment was transferred to pWS3-*msrCdwn-*T7 to create pWS3-*msrC-*pBac-T7. To compensate for the low copy number of the *gfp* integration in the chromosome, we amplified two copies of *gfp* from pREG696-*gfp* (laboratory collection) using primers with different restriction sites, oVDM1016 - oVDM1026 (EcoRI-SphI) and oVDM1018 - oVDM1027 (SphI-SmaI), and consequently ligated together after digestion with SphI. This construct with two *gfp* genes in tandem was inserted into pWS3-*msrC-*pBac-T7 via EcoRI-SmaI digestion creating the complete *msrC::gfp* template. This template was amplified using oVDM1052-oVDM1053 and transferred to pVDM1001 by digestion with SmaI creating pVDM-*msrC::gfp*.

To perform the chromosomal modifications we first transformed E745 with pVPL3004, with selection for transformants by plating on BHI with 50 μg ml^−1^ erythromycin and 24h incubation at 37°C. Presence of pVPL3004 in E745 was confirmed via PCR using primers oVDM1005 - oVDM1006. A colony positive for pVPL3004 was made competent to receive either pVDM- *ΔlacL* or pVDM-*msrC::gfp*. After transformation with these vectors the transformants were selected on BHI agar with 200 μg ml^−1^ spectinomycin and 70 μg ml^−1^ erythromycin and incubated 48-72 h at 30 °C. Successful deletion of *lacL* was confirmed by PCR with primers oVDM1007-oVDM1008. Insertion of *gfp* was confirmed by PCR with primers oVDM1009-oVDM1011.

### Curing of CRISPR and Cas9 plasmids

A colony that was positive for the desired mutation was transferred to 200 ml BHI without antibiotics and incubated overnight at 37°C at 250 rpm after which 200 μl was transferred to 200 ml pre-warmed BHI and incubated overnight at 37 °C. This process was repeated a third time after which a 100 μl sample was taken and diluted 1000 times of which 25 μl was transferred and spread on a BHI agar plate. After 24 h incubation at 37°C, 50 colonies were transferred to BHI agar, BHI agar with 200 μg ml^−1^ spectinomycin or BHI agar with 50 μg ml^−1^ erythromycin. After incubation overnight at 37°C the plates were examined for colonies that were susceptible to both spectinomycin and erythromycin. Curing of the Cas9 delivery vector pVPL3004 and the CRISPR-containing vectors derived from pVDM1001 was confirmed via colony PCR using the primer sets oVDM1054-oVDM1055 and oVDM1056-oVDM1057, respectively

### Flow cytometric analysis of GFP fluorescence in E745

To confirm the phenotype of the *gfp* integration mutant, cultures of E745, E745::*msrC::gfp*, and E745 + pREG696-*gfp* in 3 ml BHI, supplemented with 250 μg ml^−1^ spectinomycin if required, were started and incubated overnight at 37 °C. The fluorescence of the cultures was then determined by flowcytometric analysis after adjusting the cultures to an OD_600_ of 0.2. These were then diluted 25-fold in a 2-ml volume of PBS of which 200 μl was transferred to a round bottom 96-well plate, which was placed into a MACSQuant (Miltenyi Biotech) machine. Flow cytometric analysis was performed by measuring fluorescence at 488 nm excitation and 525 nm emission at 35.000 events in total. Bacteria were gated on single cells based on forward and side scatter. Data was further processed in FlowJo (FlowJo LLC).

## Results and Discussion

### Implementation of CRISPR-Cas9 mediated genome editing in *E. faecium*

We initially attempted to combine single-stranded DNA recombineering and CRISPR-Cas genome editing in *E. faecium*, as was previously demonstrated in the lactic acid bacterium *Lactobacillus reuteri* (Oh and Van Pijkeren 2014). We were, however, unsuccessful in generating mutants in *E. faecium* using this methodology. Either not enough oligonucleotides were transformed into the cells due to the inherent low transformation efficiency in *E. faecium*, or the activity of the single-stranded DNA binding protein RecT was too low to support incorporation of the oligonucleotide into the chromosome. We then decided to adapt the *L. reuteri* system by relying on the high intrinsic recombination rate of *E. faecium* for allelic exchange and by using CRISPR-Cas9 to counter select against wild-type cells. For this we used the vectors pVPL3004, which encodes Cas9, and pVPL3115, encoding the CRISPR array to which the protospacer target sequence can be added. To facilitate further adaptations needed for genomic modifications we transferred the CRISPR guide RNA section from pVPL3115 to the vector pWS3 to create pVDM1001. This plasmid has the benefit of having a temperature-sensitive replicon for Gram-positive bacteria and can replicate in *E. coli* EC1000, facilitating further cloning procedures.

The *E. faecium* CRISPR-mediated genome engineering plasmid thus relies on pVPL3004 and the novel vector pVDM1001 being present in the strain of interest (Figure 1A). The general workflow is depicted in Figure 1B. In short, pVPL3004 was first transformed into *E. faecium* E745 to allow for CRISPR-based genome modifications. We then exchanged the control protospacer in pVDM1001 for one that targets the region on the *E. faecium* chromosome that we intended to manipulate. Thirdly, we added a repair-template that contained the desired mutation. Lastly, the resulting pVDM1001-derived plasmid was transformed into E745 containing pVPL3004. Transformants were selected on BHI agar plates containing both erythromycin and spectinomycin, and were subjected to PCR to determine the recombinant genotype. As a proof-of-principle in this study, we generated a deletion mutant in *lacL* (locus tag: EfmE745_01561), the gene encoding the large sub-unit of the *E. faecium* β-galactosidase, and we integrated *gfp* in the chromosomal *msrC* gene (Singh, Malathum and Murray 2001)(locus tag: EfmE745_02638) to generate a fluorescently tagged *E. faecium* strain.

**Figure 1:**
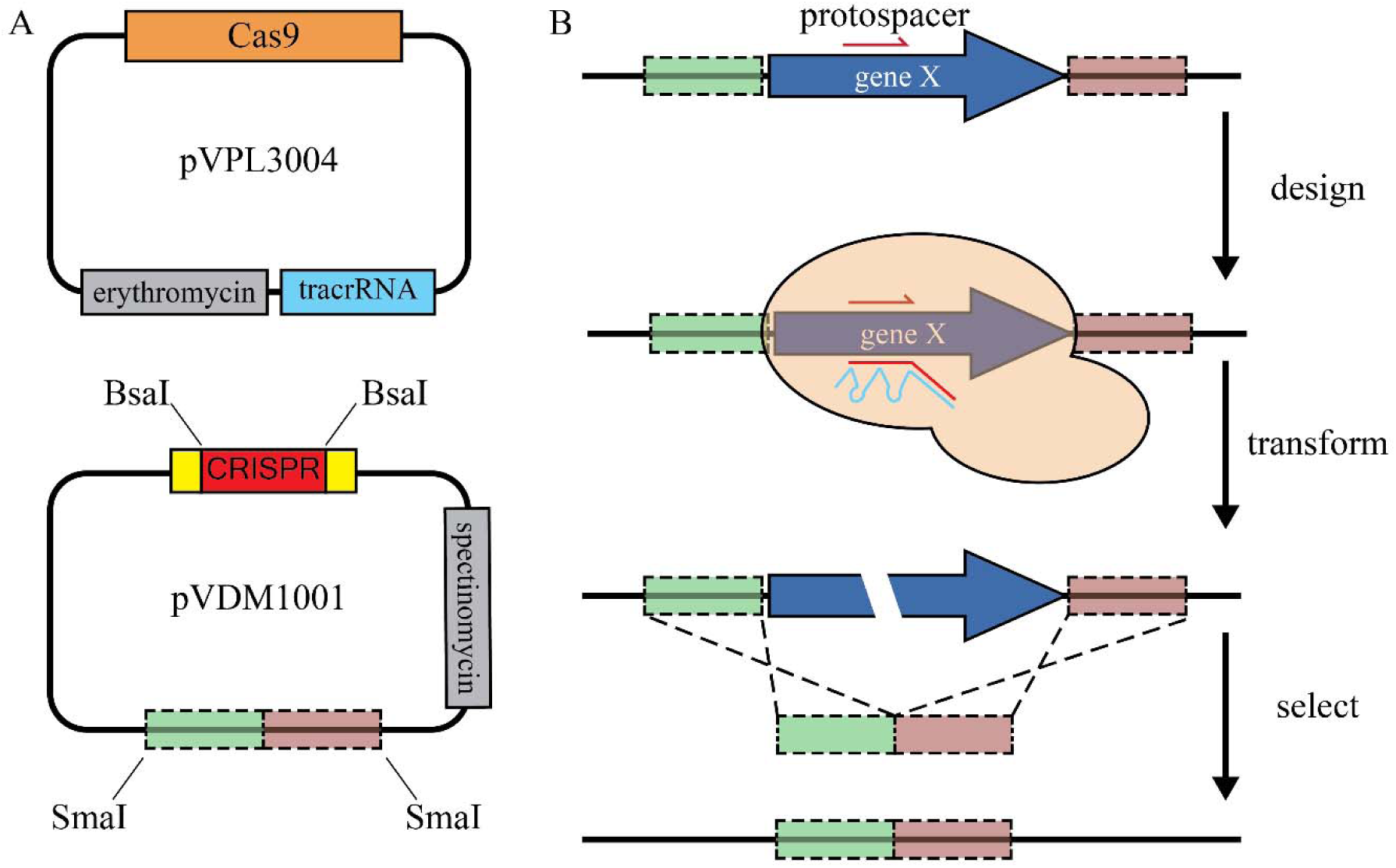
Schematic overview of the CRISRP-Cas9 mediated genome editing. This system consists of two mids (panel A), pVPL3004; which contains *cas9* from *S. pyogenes*, tracRNA and an erythromycin selection ker, and pVDM1001; which contains a CRISPR targeting the desired region, the template DNA which ies the desired mutation and a spectinomycin selection marker. The general workflow for generating mutants hown in panel B, and includes the design of the CRISPR-protospacer and repair-template which are orporated in pVDM1001. The second step is the transformation of the plasmids pVPL3004 and the relevant DM10001 derivative into *E. faecium*, followed by direct selection of the mutant.

### Generation of a deletion mutant and a chromosomal integration mutant

To delete *lacL* we adapted pVDM1001 to contain a CRISPR targeting the wild type locus of *lacL* (pVDM-x*lacL*). The vector pVDM-*ΔlacL* contained, in addition to the CRISPR targeting *lacL*, a repair template consisting of two regions flanking *lacL*, which allowed the generation of a targeted deletion mutant. To insert *gfp* in the chromosome, we created a repair template containing flanking regions of *msrC* and two copies of *gfp* in tandem as a transcriptional fusion under control of the constitutively expressed pBac promotor. We cloned the *gfp* repair template and a specific CRISPR targeting *msrC* into pVDM1001 to create pVDM-*msrC::gfp.*

In a representative experiment to generate the *lacL* deletion mutant, we transformed E745 + pVPL3004 with dH_2_O, pVDM1001, pVDM-x*lacL* (carrying a CRISPR that targets *lacL*) and pVDM-*ΔlacL* (carrying both the *lacL-*targeting CRISPR and the repair-template for generation of the *lacL* deletion mutant). This resulted in 70, 250, 68, and 80 colonies, respectively, after selection on BHI agar plates containing erythromycin and spectinomycin to select for both pVPL3004 and pVDM1001 and its derivatives. The relatively high background in the water control revealed the appearance of spontaneously erythromycin-resistant colonies. Our data also indicated that we could successfully transform pVDM1001, which lacks an *E. faecium* CRISPR-array or repair template, into *E. faecium*. The addition of a CRISPR that targets the *lacL* gene in pVDM-*xlacL* reduced colony numbers down to background levels (68 colonies versus 70 in the water control), suggesting that CRISPR-Cas9 generated lethal double-stranded DNA breaks in the *E. faecium* chromosome. Transformation of pVDM-*ΔlacL* resulted in a slight increase in colony numbers (80 colonies), potentially indicating successful integration of the repair template. This was confirmed by PCR (Figure 2A) and subsequent Sanger sequencing as we found that approximately 15% of screened colonies were *lacL* deletion mutants. We obtained comparable results in our attempt to integrate *gfp* in the *msrC* gene, with a background of spontaneously erythromycin-resistant mutants in the control experiments but a higher number of transformants upon electroporation with pVDM-*msrC*::*gfp* (data not shown). Our overall success rate in generating mutants was considerable higher in comparison to the homologous recombination-based technique we previously developed (Zhang *et al.* 2012), in which we routinely have to screen 100 or 200 colonies, after several days or even weeks of sub-culturing, before we can isolate the desired mutant that had undergone a double cross-over event.

**Figure 2:**
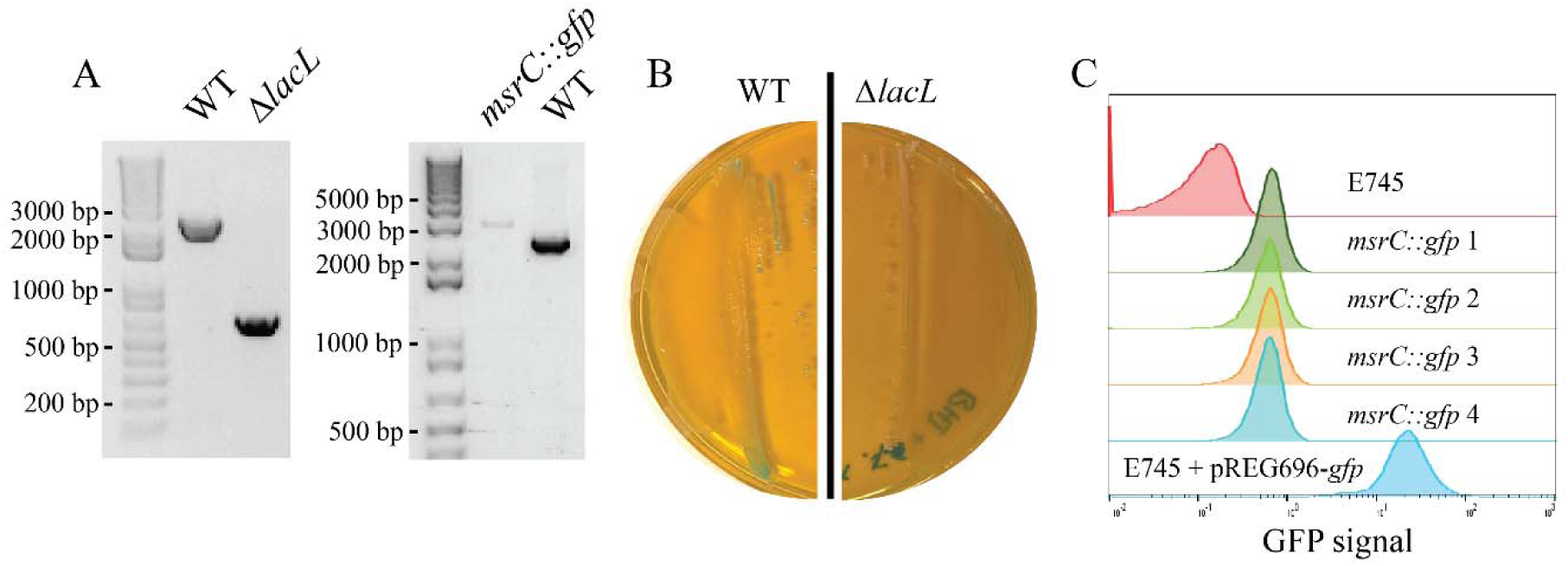
Generation and phenotypes of the δ*lacL* and *msrC::gfp* mutants. A) Confirmation of *lacL* deletion and *gfp* insertion into *msrC* via PCR. Deletion of *lacL* results in a 1800 bp reduction in size of the PCR product from 2.5 kbp to 0.7 kbp, while insertion of the *gfp* construct into the *msrC* site results in a shift from 2.8 kbp to 3.2 kbp B) Growth of wild-type E745 and δ*lacL* on BHI with 0.2% X-gal. C) Flow cytometric analysis of GFP fluorescence levels, from top to bottom, wild-type E745, four different *msrC::gfp* clones and, as a positive control, E745 containing pREG696-*gfp*.

Once we confirmed that we had successfully generated the *lacL* deletion mutant and the *msrC::gfp* insertion mutant, the CRISPR-related plasmids were cured by sub-culturing in BHI broth without antibiotics for three days, or between 20 and 25 generations. Between 50 and 100 colonies isolated from this culture were then transferred to three different BHI agar plates, i.e. BHI agar without antibiotics, BHI agar with spectinomycin and BHI agar with erythromycin to isolate colonies that had cleared both pVDL3004 and the pVDM1001-derivative. Two representative examples of experiments in which we cured the pVPL3004 and the pVDM1001-derivative are shown in Figure 3. Curing ratios for pVPL3004 were typically around 60-90% while pVDM1001-derived vectors was more difficult to cure as 1-5% of colonies had lost the vector. Typically, we obtained 3 to 5 colonies in which both plasmids had cleared per 100 colonies.

**Figure 3:**
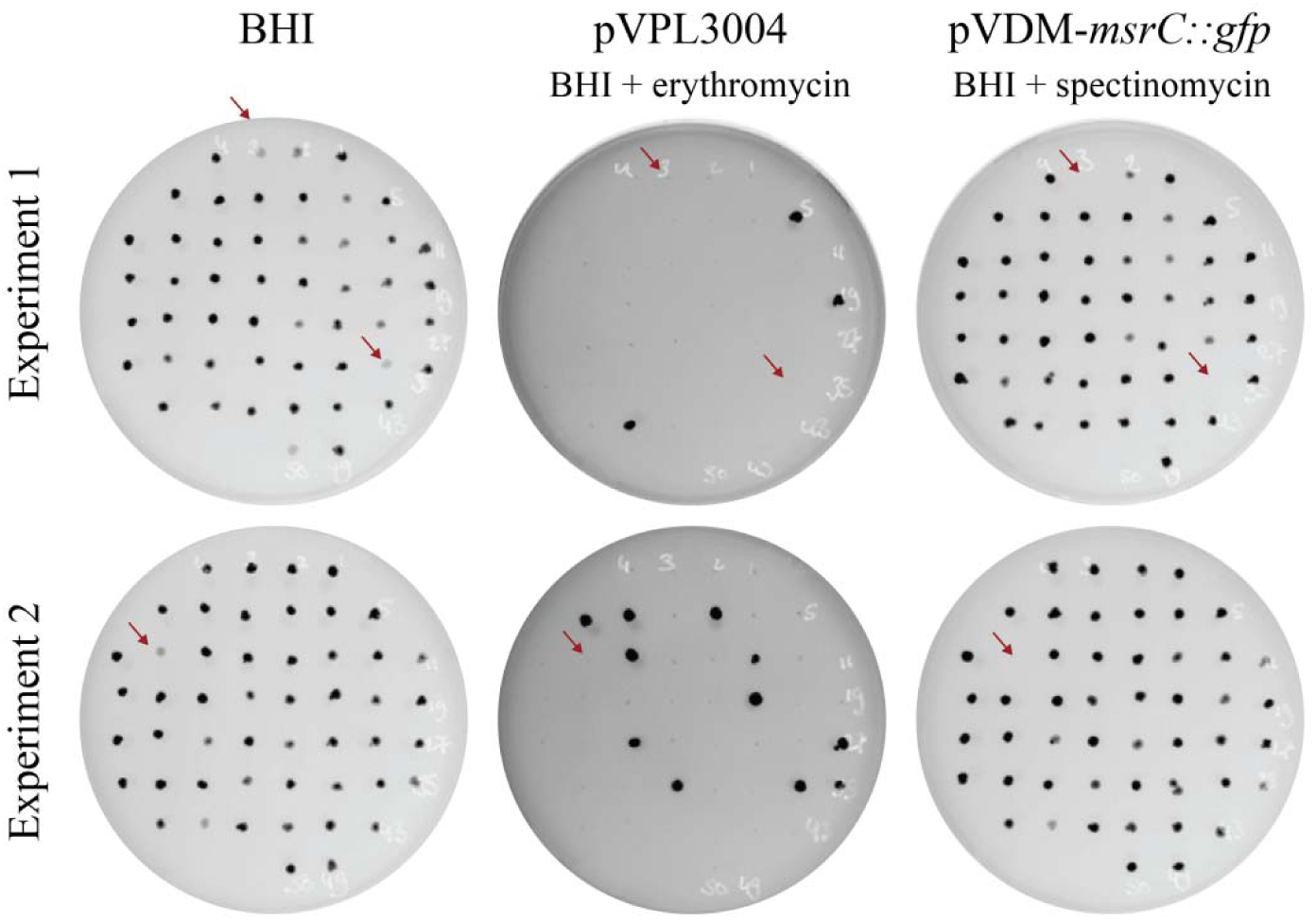
Clearing efficiency of pVPL3004 and pVDM-*msrC::gfp*. After days of sub-culturing to clear the plasmids, 50 colonies per mutant transferred to BHI, BHI + 50 µg/ml erythromycin and BHI 200 µ spectinomycin to screen for clones that have lost both plasmids (indicated b red arrows). The overall clearance of pVPL3004 is 80-90% and of pV *msrC::gfp* is 2 - 5%, resulting in at least 1 colony that has lost both plasmids results show results of two independent experiments to clear pVPL3004 pVDM-*msrC::gfp* from the insertion mutant. Colonies were visualized by ImageQuant LAS4000 imager through their production of GFP. Note tha fluorescent signal is lower in the *gfp* integration mutants than in the colonies w *gfp* is still present on a multi-copy plasmid.

### Phenotypic characterization of E745 *ΔlacL* and E745 *msrC::gfp*

Wild-type (WT) E745 and E745 *ΔlacL*, which were cleared of pVDL3004 and pVDM-δlacL as outlined above, were grown on BHI supplemented with the chromogenic substrate X-gal to confirm that the genomic alteration affected β-galactosidase activity. While WT colonies were light blue upon growth on medium containing X-gal, the E745::*ΔlacL* colonies were creamy white (Figure 2B), indicating that they could no longer convert X-gal due to the lack of an active β-galactosidase. We determined production of GFP by flow cytometry (Fig. 2C) and we found that the GFP signal is higher in E745 *msrC::gfp* compared to WT, but considerably lower than the strain in which *gfp* is carried on a plasmid. This most likely reflects differences in copy number of the chromosomally integrated *gfp* construct versus *gfp* carried on the multi-copy pREG696 plasmid.

## Conclusion

In summary, we applied CRISPR-Cas9 as a counter-selection strategy to aid in the generation of targeted modifications in the chromosome of a clinical strain of *E. faecium*. Our approach for genome editing in *E. faecium* does not require specialized media and does not leave a scar in the chromosome. Mutants could be efficiently identified by PCR and the plasmids used to generate the mutants were readily cured. In comparison with our previous protocol (Zhang *et al*., 2012), processing time was reduced by up to 2 weeks and the total number of colonies that need to be screened is reduced by approximately 4-fold. It is important to note that the use of CRISPR-Cas9 allowed us to generate deletion mutants but also to insert genes into the genome, which can be useful for a number of applications. The stable insertion of fluorescent or bioluminescent tags into the genome can be of particular use during *in vivo* experiments, e.g. to track colonization and infection by *E. faecium*. We note that the CRISPR/Cas9 system described here can be improved further, e.g. by changing the selection markers to reduce the number of spontaneously resistant colonies. Native CRISPR systems are relatively rare in multi-drug resistant clinical *E. faecium* strains (Palmer and Gilmore 2010; Lebreton *et al.* 2013) and there is therefore little risk of interference with the system we implemented here.

Even though *E. faecium* is broadly recognized as an important multi-drug resistant nosocomial pathogen, there is still a limited mechanistic understanding of its basic biology and the traits that contribute to its transition from gut commensal to opportunistic pathogen. Efficient *E. faecium* genome editing tools are essential to perform functional studies that can inform effective intervention and treatment strategies. The CRISPR-Cas9-based approach described here improves the current genetic toolbox for *E. faecium* and we anticipate that it will accelerate research into this species. We note that the approach we developed here for *E. faecium* might also be successfully implemented in other enterococci and low-GC Gram-positive bacteria with high recombination rates.

