## Supplementary material for "CRISPR-Cas9 mediated genome editing in vancomycin resistant *Enterococcus faecium*": Table S1

Supplementary Table 1: Sequence of the repair template used for the deletion of *lacL*.

*lacL* repair template

GAGTCCTTTTGAAGAAAATATTGCCGCATGGATGTTTGTGTCTCCTAAACAAGATGAAGCTATTGTGTTTTTAGGGAGGATACTAGCTTCAGCTCAACCAGCGTTCCATGAAGTATATCTGATGGGGTTAGATGATGAGGCACTTTATCAGGAACAGACCTCGAAGCGGATATTTTCGGGGGCCGAATTGATGACAGTTGGACTTTACTTCCCCGATTTTCAAGGTGATTTCCAAACAGAACTGCTTCATTTCAAAAAGTTATGAGAGAGAAGGAAAAAAGTATGAAAGCAAATATAATGATCCAATCACAGGCAGAGAAGTGATGCGCTATGGCGGTGACTTTGACGATAAACCAAGTGACTATGAATTCTCAGGGAATGGGATCGTTTTTGCAGATGGACAAGAAAAACCCGCCATGCAGGAGGTAAGATATTATTATGAAAAATACAGTAAATAAAAGTCATATGGATACGGAAAAAGTTGCAATCGTCTTCGGCGACTGTACATTAGGTGTCAAATCGGGGAATACGCATTATATTTTTTCTTATACAAGAGGCGGACTGGAATCGCTCAATAAAAACGGAAAAGAATGGCTA
